# Parental control: ecology drives plasticity in parental response to offspring signals

**DOI:** 10.1101/2021.11.18.464426

**Authors:** S.M. Caro, A.C. Velasco, T. van Mastrigt, K. van Oers, A.S. Griffin, S.A. West, C.A. Hinde

## Abstract

Different bird species have completely different parent-offspring interactions. When food is plentiful, the chicks that are begging the loudest are fed the most. When food is scarce, bird species instead feed the largest offspring. While this variation could be due to parents responding to signalling differently based on food availability, it could equally be due to offspring adjusting their behaviour, or to variation in information availability. We tested between these competing explanations experimentally, by manipulating food availability in a population of wild great tits, *Parus major*, while standardising offspring behaviour and size. We found that when food was more plentiful, parents were: (1) more likely to preferentially feed the chicks that were begging the most; and (2) less likely to preferentially feed larger chicks. In addition, we consistently found these same patterns, in a meta-analysis across 57 bird species. Overall, our results suggest that parents have more control over food distribution than offspring do, and that they flexibly adjust how they respond to both offspring signals and cues of offspring quality in response to food availability. Consequently, depending upon environmental conditions, predictably different signalling systems are favoured.

## Introduction

In species where parents care for multiple offspring at the same time, families are constantly negotiating how much parents will invest in each offspring. Yet the outcome of these negotiations is completely different in different bird species (Caro et al. 2016). At one extreme, in some species, the chicks in worse condition beg the most, and gain the most food from their parents. At the other extreme, in other species, begging appears to be ignored, and the largest chicks obtain the most food. Evolutionary theory and a comparative, across-species study have suggested that this pattern reflects parents’ adjusting their feeding strategy in response to environmental conditions and food availability (Davis et al. 1999; Kilner 2002; Mock et al. 2011; Grodzinski and Johnstone 2012; Caro et al. 2016; Koykka and Wild 2018). When food is plentiful, parents will be able to rear all their offspring, and so should preferentially feed the offspring in greatest need, which can be signalled by begging (Godfray 1995; Davis et al. 1999). In contrast, when food is scarce and only a fraction of offspring can be raised, parents should preferentially feed the best quality offspring based on size cues (Caro et al. 2016). This hypothesis posits that differences across species are determined by parents adjusting their responses to signals in response to environmental conditions.

However, this hypothesis was based on observational data, and is open to alternative explanations. First, changes in communication patterns could be controlled by either receivers (parents) or signallers (offspring) (Kilner and Hinde 2008). If offspring control resource allocation via direct competition, then the most competitive offspring should receive the most food. Consequently, in situations where competitive ability shows greater variation, which could be when food is scarce, we would expect signals of need to have less influence on food distribution (Royle et al. 2002; Parker et al. 2002). Second, parents could have access to different information under different environmental conditions, which would constrain their ability to respond to signals (Kilner and Hinde 2008). For example, if all offspring beg at maximum intensity in worse environments, parents cannot use begging to distinguish between offspring. Third, it is only known that across-species differences are correlated with environmental conditions—it is not known whether individuals are plastic in how they communicate based on local conditions. We might only expect plasticity to evolve in species that experience variable ecological conditions within or between breeding bouts (Forsman 2015). If parents could flexibly vary their decision rules according to the environment, that would substantially increase the level of control that parents have within the family (Kilner and Hinde 2008).

We used an experimental approach to directly test whether parents respond differently to begging depending upon food availability in great tits, *Parus major*. This species lives in temperate regions and is exposed to variable breeding conditions across years, and could therefore be expected to evolve plasticity in response to offspring signals and cues. To distinguish between parental and offspring responses to food availability, we both provided supplemental food to some parents and cross-fostered offspring before observing behaviour. This allowed us to observe parents that had and had not been supplemented, interacting with foster broods that were similar in begging intensity, chick size cues, chick competitive asymmetry and supplementation history, as foster broods comprised half supplemented, half unsupplemented chicks. Next, to investigate whether plasticity is a general pattern across species, we conducted a phylogenetic meta-analysis, examining the pattern of behaviour within 57 bird species. We assessed within-species plasticity by quantifying each species’ change in responsiveness to begging or chick size as environmental quality improves.

## Materials and Methods

### Experimental study

#### Study area and species

Great tits (*Parus major*) are a common passerine bird distributed across Eurasia. They are primarily insectivorous while feeding young, with highly variable food availability both geographically and temporally (van Balen 1973). This variation in ecological conditions within and between breeding bouts makes great tits a prime candidate for studying the evolution of flexibility in parental provisioning strategies. We studied a wild population of great tits living in a mixed pine-deciduous forest (Boslust) covering approximately 75 ha in The Netherlands (5°85’E, 52°01’N). From March through June 2017, we monitored 130 nest boxes, and were able to include 34 broods in our study. We checked nest boxes every other day to determine the onset of egg laying and clutch size. We began visiting nests daily the day before hatching was expected to determine hatch date (day 0), brood size and mortality rates. Mean clutch size was 9.29 ± 0.23 SE eggs, and mean brood size at hatching was 8.82 ± 0.26 SE in our study population. All of the study broods hatched within 9 days of each other. Across all broods, 10.9% of chicks (33 of 302 chicks) died in the first week after hatching.

#### Experimental procedures

In order to simulate variation in ecological conditions, we experimentally manipulated food availability in an alternating pattern: half of the broods received supplemental food (mealworms and wax worms), while the other half experienced natural conditions (see supplemental methods for details).

We wanted all parents to be exposed to equivalent information from their broods during filming, so that we could rule out the possibility that offspring are driving any differences in parental provisioning preferences. We therefore standardized brood size and offspring supplementation history across all broods immediately before filming. We cross-fostered chicks on the filming day (8 days after hatching) to create experimental filming broods of 7 (27 broods) or 6 chicks (4 broods). Approximately half of the chicks in each filming brood came from a supplemented nest, and the other half of the chicks came from an unsupplemented nest. Fostered chicks were the same age as the parents’ biological brood. Parents were not filmed with their own chicks. We also wanted to ensure that there would be sufficient and equivalent variation in offspring size so that parents could use this information during food allocation. To create an even distribution of weight and prior weight ranks in filming broods, we ranked chicks by weight in their biological nests. We assigned the heaviest chick to filming brood A, and the second heaviest to filming brood B, the third heaviest to brood A, etc. We alternated this pattern at each nest. We wanted to ensure that there would be enough variation in begging intensity and to ensure initial begging intensity would vary across weight ranks. We hand-fed half of the chicks in each filming brood to satiation, in an alternating pattern by weight rank (see supplemental methods for details). This ensured that not all chicks begged maximally during filming and that not all small chicks begged at highest intensity the whole time.

Thus, parents in both treatments were filmed at the same time of day, feeding broods of an equivalent size and begging intensity, comprising unrelated supplemented and unsupplemented chicks, half of which were satiated when the filming began.

#### Details for great tit experimental supplementation

To ensure only experimental broods received extra food, and to avoid changes to nest defense associated with positioning the food near the nest box, we installed a small feeding tray inside each nest box (Verhulst 1994; Grieco 2003; Eeva et al. 2009). This was done during incubation at all nests. No broods were deserted after the introduction of the tray. Each day for the first week after hatching, we provided a *c.* 20g mixture of live meal worms (*Tenebrio molitor*) and rehydrated wax worm larvae (*Galleria mellonella*) cut into 0.25 cm pieces to supplemented nests. This represents approximately 20% of the daily nutritional needs of the brood (van Balen 1973; Eeva et al. 2009). We checked whether great tits were using the food by placing cameras into 2 nests during the supplementation period. We observed parents taking food from the trays and directly feeding their offspring (Supplementary Movie 2), and parents also ate the food themselves. Either outcome serves to increase environmental conditions for the parents. Control nests were also visited each day so that all nests received comparable experimental disturbance, and an empty tray was placed in the nest box.

We alternated experimental treatments by assigning the first brood of the day that had hatchlings to the supplemented treatment, and then the next brood the unsupplemented (control) treatment. We reversed this order each day. We did not pre-randomize because we wanted to equalise hatch date within each treatment. Supplemented and unsupplemented nests varied slightly in clutch size (supplemented 9.81 +/− 0.33se, unsupplemented 8.82 +/− 0.32, p = 0.038*), but not in brood size (supplemented 9.18 +/− 0.36se, unsupplemented 8.59 +/− 0.36, p = 0.26) or hatch date (supplemented 25.29 +/− 0.58se, unsupplemented 25.18 +/− 0.57, p = 0.89). The difference in clutch size was driven by one unsupplemented nest with only 6 eggs; removing this nest or including clutch size as a control variable did not change the results of our parental response model.

#### Details of cross-fostering and hand-feeding

All cross-fostering was done in the morning as soon as possible prior to filming, and all filming occurred between 7:00 and 15:00 (83% of feeding visits occurred between 9:00 and 13:00).

Hand-feeding protocol: We ranked chicks by weight in their filming nests. We assigned chicks to be handfed or not handfed in an alternating pattern by weight rank, which was reversed at each nest. For example, in filming brood A, the heaviest chick was handfed and the second heaviest was not, while in filming brood B the heaviest chick was not handfed. Immediately prior to filming, we hand-fed chicks in an artificial nest containing a cloth wrapped hand-warmer. We fed the selected chicks with Nutribird A 19 high energy bird food using a 5 mL syringe. We continued feeding until begging had ceased and could no longer be induced by whistling and tapping the sides of the bill with a syringe, indicating the chicks were probably satiated, as in (Kilner and Davies 1998).

#### Video data

We filmed parents feeding their foster broods 8 days after hatching (see Supplementary Movie 1 for an example of video data). We installed an infrared camera inside the lid of a nest box the day prior to filming in order to habituate parents. We paint-coded all chicks with a dot of red, non-toxic acrylic paint on the head (Kate Lessells, pers. comm.) just prior to filming, so that we could individually identify chicks in the videos. We excluded the first 30 minutes of filming to ensure that parental and chick behaviour had enough time to return to normal after cross-fostering and to give us enough time to leave the area. We did not provide supplemental food to the parents on the filming day.

All videos were coded by the same observer, blind to the experimental treatment and to chick weight ranks. The order in which the observer coded the videos was random with respect to whether nests were supplemented and unsupplemented. Adult identity was determined by the difference in crown feather glossiness of males and females, and confirmed by nest cleaning behaviour, which only females perform (Christe et al. 1996). For each feeding visit, the observer recorded the sex of parent, the identity of the fed chick, and the begging intensity of all chicks. The observer recorded 20 feeding visits per parent or 4 hours of filming, whichever came first.

#### Begging intensity

Begging intensity was coded on a standard scale, following Hinde 2009, adapted from Kilner (1995): *0 = non-gaping, 1 = gaping with a bent neck, 2 = gaping with neck stretched out, 3 = gaping with raised body* (Kilner 1995; Hinde et al. 2009). We quantified relative begging intensity by dividing the begging posture of each chick by the mean posture of all begging chicks on that feeding visit. Values greater than 1 indicate that parents preferentially fed chicks with a higher posture score than their nest mates. This relative measure accounts for differences in overall begging intensity on different feeding visits, which could confound measures of food distribution based on absolute begging intensity (Hinde et al. 2009).

#### Chick size

We ranked chicks by weight in their filming brood, with chick 1 being the heaviest and chick 7 being the lightest. Using weight rank as opposed to absolute weight makes nests more directly comparable—parents may always prefer feeding the largest chick, whether the largest chick weighs 12g or 10g. A priori, we assumed that weight rank need not have a linear effect. Parents may treat large and medium sized chicks differently than they treat small chicks, since the smallest chicks may be most vulnerable to starvation (Magrath 1990; Forbes et al. 1997; Amundsen and Slagsvold 1998; Theofanellis et al. 2008; Podlas et al. 2013). Furthermore, the difference in absolute weight between ranks was lower in the middle of hierarchy (0.68g) than at either end (1.05g). We therefore included the quadratic term for weight rank in analyses.

#### Statistical analysis

We checked whether our supplementation treatment was biologically relevant by investigating its effect on the likelihood of brood reduction (whether at least one chick died) in the first week after hatching before any cross-fostering took place. The effect of supplementation on likelihood of brood reduction likelihood was assessed using a binomial linear model in lme4 in R, while controlling for clutch size, brood size, and hatch date (Bates et al. 2015). The extent of brood reduction was assessed using a quasi-poisson linear model to account for zero-inflated count data, while controlling for clutch size, brood size, and hatch date (Bates et al. 2015). Chick mass one week after hatching was assessed using a linear mixed model, while controlling for clutch size, hatch date, and nest ID (Bates et al. 2015). We standardised and centered control variables (Cohen et al. 2003).

We analysed parental provisioning using a Bayesian logistic mixed model (MCMCglmm) in R (Hadfield 2010; R Core Team 2013). We used uninformative priors, ran the model for 700,000 iterations with a burn-in of 150,000 and a thinning interval of 10. We assessed the convergence of the MCMCglmm model by visual inspection of convergence plots and geweke plots (Hadfield 2010; 2012). The response variable was whether a chick was fed or not. We included nest ID, parent ID, chick ID, and feeding visit ID as random effects. We analysed a three-way interaction between supplementation treatment, relative begging intensity, and weight rank as the fixed effect. We were interested in this interaction because our main hypothesis was about the moderating effect of supplementation, and because parents may respond differently to begging of different offspring (van Heezik and Seddon 1996). We tested whether weight rank had a non-linear effect. We included both the linear and quadratic 3-way interaction terms in the same model, along with all possible 2-way interactions using both the linear and quadratic terms (Ganzach 1997). If the quadratic interactions were not significant, we would have removed them. Including polynomial terms in interactions can lead to false positives or negatives due to collinearity (Ganzach 1997). Centering variables reduces this collinearity between polynomials, and so we scaled and centred begging and weight rank (Cohen et al. 2003; Dalal and Zickar 2011).

Of the 34 broods filmed for our study, we excluded one brood because the parents abandoned during filming, and two broods that had fewer than 20 feeding visits. We excluded data from four parents with fewer than 15 observed feeding visits. We excluded six feeding visits where the begging posture of more than two chicks was unknown. Our final sample size for the analysis of parental provisioning was 14 supplemented nests, 15 unsupplemented nests (54 adults, 199 chicks, 1121 feeding visits). We analysed the full data set as well, and there were no qualitative differences in the results or in their statistical significance.

### Comparative study

#### Data collection for the meta-analysis

To determine whether birds show a consistent adjustment of feeding rules based on local conditions across species, we collected data on within-species changes in the strength of the relationship (correlation coefficient) between feeding and begging in 17 species, and feeding and size cues in 52 species (719 effect sizes from 145 studies; Data S1; Fig. S1). We conducted a literature search on Web of Science and Google Scholar using the keywords ‘beg’, ‘parent-offspring’, ‘bird’, ‘begging’, ‘communication’ and ‘provision’ (see Fig. S1 for PRISMA flowchart detailing data collection and exclusion criteria). We included all papers with any measure relating to the relationship between parental food allocation and offspring behavioural begging or size cues. We excluded species that did not have data on these relationships in more than one environment condition, since we were interested in the change in the strength of these relationships over different ecological conditions. We excluded studies if it was impossible to determine whether parents were responding to begging or to size cues. We excluded studies where offspring signals were structural (such as mouth colour), rather than behavioural (such as begging postures), as these may represent different signalling systems (Caro et al. 2016). There was not enough within-species data on structural signals in different environments include them. We only included effect sizes for the relationship of begging on within-brood food allocation, rather than on increases in overall parental feeding effort, as these represent fundamentally different aspects of parental care. We excluded data on species that lay only one egg per brood, as selective pressures on these species are likely to differ from species laying multiple eggs per brood. If relevant data were given in papers without statistical tests, such as raw means and standard errors, we estimated effect sizes (Borenstein et al. 2011). This resulted in a dataset of 719 effect sizes from 145 studies on 57 species (Data S1).

### Environmental quality

We categorized populations as experiencing normal, better than normal, or worse than normal environments, based on experimental manipulations of long-term chick condition (parents were fed reduced or supplemented diets, or chick demand was artificially increased or decreased), ecological measures (such as prey density, date or rainfall), or average offspring mortality across different years in long-term observational studies (Caro et al. 2016). These measures were not always directly related to food availability, but they likely captured variation in some ecological aspect relevant to offspring condition. If no information on environmental quality was available, data were classified as normal conditions.

#### Measures of feeding, begging and size cues

Many aspects of behavioural begging were reported in the literature, such as begging amplitude, duration, latency, likelihood, call structure, and begging postures. Different measures of food allocation were also presented, such as weight gain over a short time period, actual food intake, number of food items received, likelihood of being fed, growth rate, and mortality. We assumed all measures of begging intensity and feeding preferences were functionally equivalent, and so included all reported statistics in our analyses. Because test statistics were converted to a standardized scale, differences between the various measures of begging intensity or feeding preferences should not influence the overall trends seen. A previous comparative analysis, using a more comprehensive dataset, found no impact of study methodology, such as which measure of feeding preference was used or whether studies were experimental or observational, on the effect size of begging or size cues on feeding preferences (Caro et al. 2016).

#### Statistical analyses

We transformed any test statistic measuring either an effect of begging or size cues on feeding into a standardized effect size (Fisher’s Z-transformed correlation coefficient) (Borenstein et al. 2011; Grissom and Kim 2011; Koricheva et al. 2013). These correlation coefficients follow a normal distribution, account for different scales in their original measurements, are well suited to the ordered nature of the data, and are more straightforward to interpret than standardized differences in means (Borenstein et al. 2011; Grissom and Kim 2011; Koricheva et al. 2013). Fisher’s Z-transformed correlation coefficients were analysed using the MCMCglmm package in R, which implements Bayesian generalized linear mixed models with Markov chain Monte Carlo methods (Hadfield 2010; R Core Team 2013). We ran bivariate response models so we could measure phylogenetic covariance between responsiveness to begging and responsiveness to size cues. We weighted models by sample size, and controlled for phylogeny and repeated measures on the same study and species. Our measure of sample size was the number of broods used to generate the original test statistic, because this is a standard measure across studies, and conservatively avoids pseudoreplication if chick number or number of observations were used as the sample size. We treated environmental quality as a three-level ordered categorical variable. To control for phylogeny, we obtained phylogenies from Birdtree.org, ran models on 4 random phylogenetic trees with Ericson and Hackett backbones, and then averaged model results (Jetz et al. 2014).

## Results

### Great tit experiment

We found that parental provisioning rules were flexible in response to environmental conditions – parents responded differently to offspring begging and size, depending upon whether parents had received supplemental food in the previous week (interaction between supplementation, quadratic weight rank and relative begging intensity: 95% CI −1.09 to −0.03, pMCMC = 0.041*; Fig. 1; Table S2). In all nests and for all chicks, the likelihood of being fed increased with higher relative begging intensity (95% CI 1.91 to 2.87, pMCMC < 0.0001***; Table S2), but chick size mediated how much begging increased the likelihood of being fed. In nests that had been supplemented (Fig. 1a), parents responded primarily to begging signals, responding equally to the begging of all chicks regardless of size. However, in unsupplemented nests (Fig. 1b), parents responded more to size by responding more to the begging of larger chicks than to the begging of the smallest chicks.

**Fig. 1.**
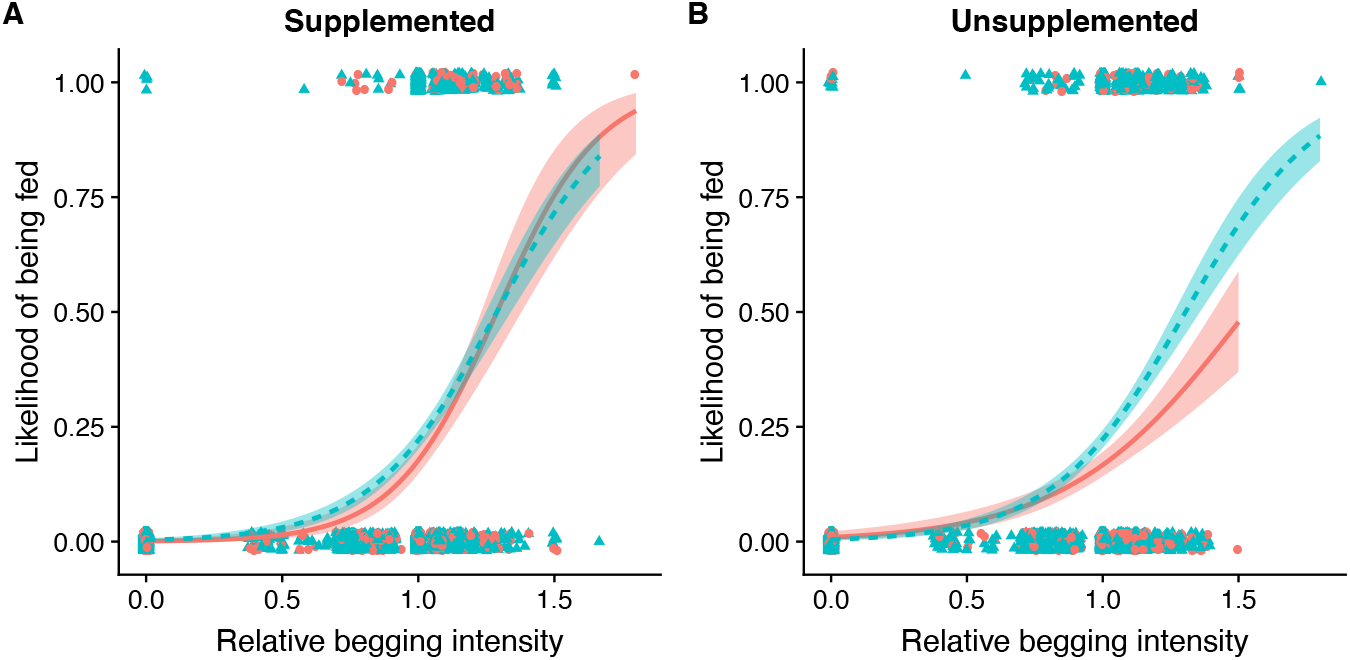
A chick’s likelihood of being fed depends on its relative begging intensity, size rank, and whether its parents were supplemented. In supplemented nests (a), higher relative begging intensity led to a greater likelihood of being fed for all chicks equally. In unsupplemented nests (b), smaller chicks showed less of an increase in their likelihood of being fed based on begging intensity than larger chicks did (95% CI of the interaction = −1.09 to −0.03, MCMCglmm). Weight ranks 1-5 are shown in blue, while the smallest two chicks in the nest (ranks 6-7) are shown in red. We show weight categories for graphical clarity; statistical analyses report the non-linear effect of weight rank as a continuous variable. A relative begging intensity of 1 indicates a chick is begging the same as its nest mates on average, while >1 means it begged at a higher intensity. Each data point is one chick in a feeding visit, vertically jittered to show overlapping points (n = 14 supplemented nests, 15 unsupplemented nests, 1121 feeding visits).

We confirmed that this difference in food allocation could not be explained by differences in offspring behaviour or size cues. Since we swapped chicks between nests directly before filming, there was no difference in relative begging posture or in chick weights between supplemented and unsupplemented foster broods (mean of begging: t_27_ = 0.63, p = 0.53; SD of begging: t_27_ = 0.15, p = 0.88; mean of weight: t_27_ = −0.90, p = 0.38; SD of weight: t_27_ = −0.64, p = 0.53). Since we handfed a subset of chicks across the weight hierarchy, relative begging posture also did not vary by chick weight rank (t_167.8_ = 0.40, p = 0.69) or quadratic weight rank (t_174.2_ = −0.62, p = 0.54). We also confirmed that our supplemental feeding was successful in improving environmental conditions: 59% of supplemented nests had no brood reduction in the first week after hatching, compared to only 18% of unsupplemented control nests (z = 2.94, p = 0.0033**; n = 34 nests; Table S1). The total number of chicks that died per nest was also lower in supplemented nests (z = 2.10, p = 0.045*; Table S1). Chick mass of the surviving chicks on day 7 was not affected by supplementation (z = 1.58, p = 0.12; Table S1).

### Comparative study

Next, to explore whether plasticity is a general trend across birds, we conducted a phylogenetic meta-analysis on 57 bird species. Species were included if they had data on parental responsiveness to begging or chick size in multiple environmental conditions (poor, normal or good). We quantified responsiveness as the correlation coefficient (effect size) between feeding and (a) begging or (b) size. These two coefficients vary between +/− 1, with higher values indicating that either (a) begging or (b) size has a stronger effect on the likelihood that a chick is fed. We estimated plasticity by calculating each species’ change in responsiveness as environmental quality varies, i.e. the within-species slope of correlation coefficients over environmental quality. A positive slope would indicate that parents become more responsive in better environments, a slope of 0 would indicate no plasticity based on the environment, and a negative slope would indicate that parents become less responsive in better environments. If other species adjust their behaviour in the way that we have observed with great tits, then we would observe a consistent within-species pattern, with parents becoming more responsive to begging (positive slopes), and less responsive to size (negative slopes), in better environments.

We found this predicted pattern, with parents became more responsive to begging, and less responsive to size, in better environments. Specifically, we found a stronger correlation between begging and feeding in better environmental conditions (95% CI of the slope = 0.13 to 0.67, pMCMC=0.0037; Fig. 2a). Across 17 species with available data on responsiveness to begging in more than one environmental condition, 14 species showed the predicted increase in correlation strength (82%), and three species showed a decrease (18%).

**Fig. 2.**
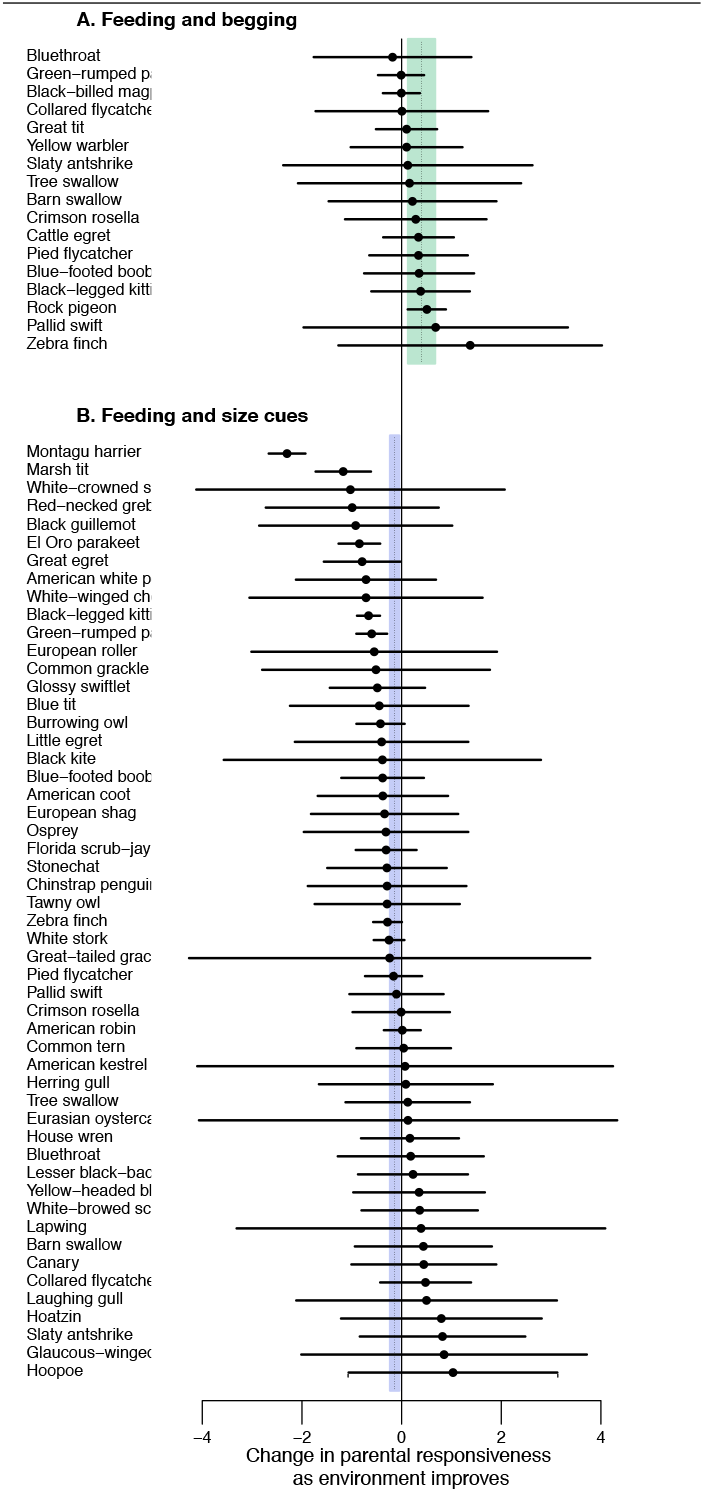
Environmental quality and parental response to offspring begging and size. Circles represent each species’ slope for the correlation coefficient between feeding and (**a)** begging, and (**b)** size cues, over environmental quality. Lines represent the 95% CI of the slope within each species. The shaded region shows the 95% CI across all species, controlling for phylogeny and weighted by sample size. Positive values indicate that parents respond more in better environments (green). Negative values indicate parents respond less in better environments (purple). Species respond more to begging (95% CI = 0.13 to 0.67, MCMCglmm, n=17 species), and less to size cues (95% CI = −0.23 to −0.05, MCMCglmm, n=52 species), in better environments. This pattern is more consistent for changes in responsiveness to begging, but is also significant for changes in responsiveness to size.

Also, as predicted, we found the opposite pattern in parental response to size cues. Within species, there was a weaker correlation between chick size and feeding in better environmental conditions (95% CI of the slope = −0.23 to −0.05, pMCMC = 0.0016; Fig. 2b). Across 52 species with available data on responsiveness to size cues in more than one environmental condition, 32 species showed the predicted decrease in correlation strength (62%), and 20 had an increase (38%). There was no species-level correlation between how responsive parents are to begging and how responsive they are to size cues (95% CI −0.54 to 0.52). These results suggest that the pattern we observed in great tits, where parents facultatively adjusted their responsiveness in reaction to local conditions, occurs consistently across a range of different bird species.

## Discussion

Our experimental and comparative results show that parents conditionally adjust how they respond to signalling, depending upon environmental conditions (food availability). Parents did not simply feed the chicks that were the largest or that begged the most. Instead, they have evolved to adjust their sensitivity to multiple sources of information depending on local conditions, in a sophisticated manner. When food is more plentiful, great tit parents respond equitably to all their offspring’s begging, but when food is more scarce, parents selectively respond more to the begging of larger chicks (Fig. 1a,b). Likewise in our meta-analysis, we found the same consistent pattern, across 57 species (Fig. 2a,b). These results show how variation in environmental quality can lead to different forms of communication, even within species.

The degree to which parents actively control food allocation as opposed to passively respond to the greatest stimulus or cede to the winner of sibling competition has been contentious (Clutton-Brock 1991; Kacelnik et al. 1995; Parker et al. 2002; Heeb et al. 2003; Ploger and Medeiros 2004). Since our cross-fostering experiment ensured there was minimal variation in brood size, competitive asymmetry and begging behaviour, changes in allocation patterns can definitively be attributed to changes in great tit parental response strategies, rather than differences in offspring behaviour or information constraints. It should be noted, however, that we examined provisioning at one point midway through the nestling period. It is possible that older chicks may be able to exert more control via scramble competition. Furthermore, given that species differed in the degree of their plasticity (Fig. 2a,b), it is probable that species vary in the actual balance of power between parents and offspring, so that in some species offspring behaviour drives changes in food distribution patterns. It may be also that some species flexibly determine their investment strategies in other ways and at other times; for example, 1) during incubation by varying the amount of hatching asynchrony (e.g. blackbirds (Magrath 1992) and European rollers (Parejo et al. 2015)); 2) during different points in the breeding season by varying how parents respond to UV signals (e.g. alpine swifts and European starlings (Bize et al. 2006)); or 3) during different times within a single breeding attempt by varying aggression towards offspring (e.g. American coots (Shizuka and Lyon 2012)). Recent work on genetic covariance and plasticity in canaries found that both offspring and parental signalling strategies varied plastically across different hunger levels (Fresneau and Müller 2019). This indicates that even if parents are plastic in their behavior and control provisioning, they may still be influenced by changes in their offspring’s behavior. Future research could continue disentangling what is driven by parental preference, by parents’ reactions to offspring signals, or by offspring directly.

What explains diversity in signalling systems is a fundamental question in signalling theory. Our results suggest that receivers control the outcome of parent-offspring communication and assess multiple sources of information from signallers. This is analogous to how females respond to multiple signals of quality in sexual signalling (Bro-Jørgensen 2010), and may be similar to aggressive signalling and other forms of communication as well. Our results highlight the need for dynamic signalling models that allow for flexibility in responsiveness based on environmental conditions, and that incorporate multiple signals and cues (Mangel and Clark 1988; Wild 2011).

## Notes

### Competing Interest Statement

The authors have declared no competing interest.

