## Supplemental Information for "Parental control: ecology drives plasticity in parental response to offspring signals"

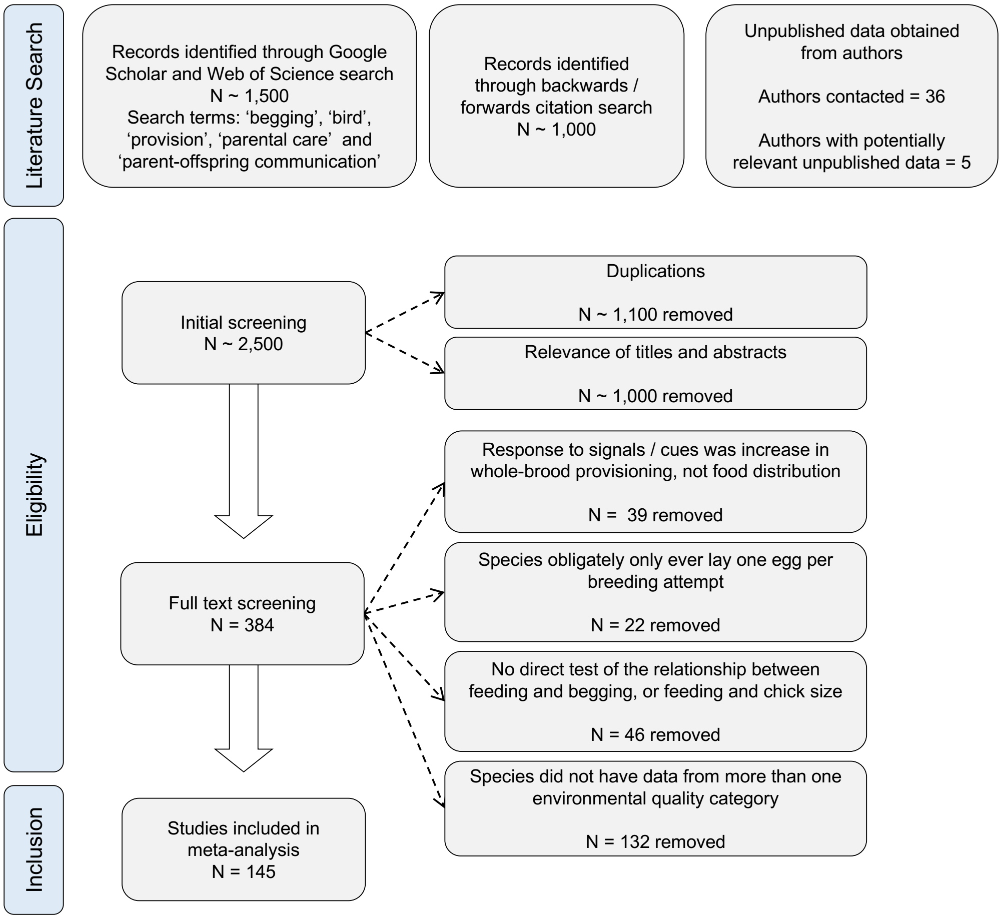


**Supplementary Fig. 1.** PRISMA flow chart of search results and the study selection process.

### Supplementary Results

**Supplementary Table 1.** The impact of parental food supplementation on offspring survival and mass

**A. Likelihood of brood reduction (yes/no)**

| | Z score | P value |
|---|---|---|
| Supplementation | 2.94 | 0.0033** |
| Clutch size | 1.94 | 0.053 |
| Brood size | -0.96 | 0.34 |
| Hatch date | -1.02 | 0.31 |

**B. Extent of brood reduction (number of dead chicks)**

| | Z score | P value |
|---|---|---|
| Supplementation | 2.10 | 0.045* |
| Clutch size | 1.76 | 0.09 |
| Brood size | 1.20 | 0.23 |
| Hatch date | -0.42 | 0.68 |

**C. Chick mass on day 7 (surviving chicks only)**

| | Z score | P value |
|---|---|---|
| Supplementation | 1.58 | 0.12 |
| Clutch size | -0.99 | 0.33 |
| Hatch date | -0.67 | 0.51 |

We tested the impact of supplementation on (**A**) the likelihood of brood reduction using a binomial linear model; (**B**) the extent of brood reduction using a quasi-poisson linear model; and (**C**) chick mass using a linear mixed model with nest ID as a random effect. N = 34 nests (17 supplemented, 17 unsupplemented); 302 chicks (154 supplemented; 148 unsupplemented).

**Supplementary Table 2. The effect of supplementation treatment, begging and size on chick feeding.**

| | **Estimate** | **95% CI** | **pMCMC** |
|---|---|---|---|
| Supplementation | -0.07 | -3.74 to -2.74 | <0.001*** |
| Weight rank | -0.24 | -0.62 to 0.14 | 0.84 |
| Weight rank$^2$ | -0.47 | -0.88 to -0.07 | 0.021* |
| Relative begging posture | 2.38 | 1.91 to 2.87 | <0.001*** |
| Supplementation: weight rank | 0.22 | -0.22 to 0.69 | 0.35 |
| Supplementation: weight rank$^2$ | 0.50 | 0.01 to 0.99 | 0.045* |
| Supplementation: begging posture | -0.04 | -0.71 to 0.58 | 0.91 |
| Relative begging posture : Weight rank | 0.12 | -0.28 to 0.54 | 0.58 |
| Relative begging posture : Weight rank$^2$ | 0.40 | -0.04 to 0.85 | 0.07 |
| Supplementation: Begging posture : Weight rank | -0.36 | -0.86 to 0.12 | 0.14 |
| Supplementation: Begging posture : Weight rank$^2$ | -0.56 | -1.09 to -0.03 | 0.041* |

MCMCglmm logistic regression on the likelihood of being fed (yes/no). Supplementation treatment was either control or supplemented. Relative begging posture is the posture of the focal chick divided by the mean posture of all begging chicks on that feeding visit. The non-linear effect of weight rank was analysed using the quadratic term. Nest, parent ID, chick ID, and feeding visit were included as random effects. N = 29 nests, 54 adults, 199 chicks, 1121 feeding visits.

**Supplementary Video Legends**

**Supplementary Video 1:** Example of a great tit feeding visit during experiment

**Supplemetary Video 2:** Example of great tit parents using the supplementary food
